# DUSP7 Regulates the Activity of ERK2 to Promote Proper Chromosome Alignment During Cell Division

**DOI:** 10.1101/2020.06.05.137364

**Authors:** Xiao Guo, Yenni A. Garcia, Ivan Ramirez, Erick F. Velasquez, Lucy W. Gao, Ankur A. Gholkar, Julian P. Whitelegge, Bobby Tofig, Robert Damoiseaux, Jorge Z. Torres

## Abstract

Human cell division is a highly regulated process that relies on the accurate capture and movement of chromosomes to the metaphase plate. Errors in the fidelity of chromosome congression and alignment can lead to improper chromosome segregation, which is correlated with aneuploidy and tumorigenesis. Here we show that the dual specificity phosphatase DUSP7 is important for regulating chromosome alignment. DUSP7 bound to ERK2 and regulated the abundance of active phospho-ERK2 through its phosphatase activity. Overexpression of DUSP7, but not catalytic dead mutants, led to a marked decrease in phopho-ERK2 and mitotic chromosome misalignment, while knockdown of DUSP7 also led to defective chromosome congression that resulted in a prolonged mitosis. Consistently, chemical inhibition of the MEK kinase that phosphorylates ERK2 or ERK2 itself led to chromosome alignment defects. Our results support a model where MEK phosphorylation and DUSP7 dephosphorylation regulate the levels of active phospho-ERK2 to promote proper cell division.

## INTRODUCTION

Critical to the fidelity of cell division is the accurate movement and alignment of chromosomes at the metaphase plate and their subsequent segregation during anaphase. Errors that compromise the precision of mitotic chromosomal congression and alignment have been linked to human developmental disorders and tumorigenesis (Gordon et al., 2012). Decades of research have highlighted the importance of protein phosphorylation as a molecular switch to regulate the activity of key proteins that are critical to cell division (Heim et al., 2017). This has been highlighted by the growing list of essential mitotic kinases and their broad array of substrates that carry out critical functions related to centrosome homeostasis, bipolar spindle assembly, kinetochore-microtubule attachments, chromosome congression, and chromosome segregation (Combes et al., 2017; Magnaghi-Jaulin et al., 2019; Saurin, 2018). A prime example is the multi-component spindle assembly checkpoint (SAC), which is comprised of several mitotic kinases including MPS1, BUB1, and BUBR1 (Sudakin et al., 2001). The SAC is activated in response to unattached kinetochores and is silenced by the achievement of the end-on interactions between kinetochores and spindle fibers, which is critical for proper chromosome segregation (Etemad et al., 2015; Foley and Kapoor, 2013; Musacchio and Salmon, 2007; Tauchman et al., 2015). Beyond well-established mitotic kinases, other less studied phospho signaling pathways have been implicated in cell division including the Wnt, mTOR and MAPK/ERK pathways, among which the MAPK/ERK pathway is phosphorylated by MEKs (mitogen-activated protein kinase or extracellular signal-regulated kinase kinase) to regulate downstream transcription factors (Bryja et al., 2017; Cuyas et al., 2014; Eblen, 2018). In *Xenopus laevis* ERK2 (extracellular signal-regulated kinase 2) is important for the spindle assembly checkpoint (Minshull et al., 1994; Takenaka et al., 1997; Wang et al., 1997). In mammalian cells ERK1/2 activity is necessary for the G1/S transition and early G2 events for the timely entry into mitosis (Meloche and Pouyssegur, 2007; Shinohara et al., 2006). However, whether human ERK2 is active in mitosis and what roles ERK2 plays in human somatic cell division remains ambiguous and controversial.

Our recent genetic RNAi screen of the druggable genome for novel factors important for cell division identified the dual specificity phosphatase 7 (DUSP7/MKP-X) (Torres lab unpublished). DUSP7 along with DUSP6/MKP-3 and DUSP9/MKP-4 are members of the cytoplasmic extracellular signal-regulated kinase (ERK)-specific mitogen-activated protein kinase phosphatases (MKPs) subfamily that share similar amino acid sequences, subcellular localizations and substrate preferences (Kim et al., 2016; Patterson et al., 2009; Seternes et al., 2019). Together the MKPs, slingshot phosphatases, PRLs (phosphatases of regenerating liver), Cdc14 phosphatases, PTEN-like and myotubularin phosphatases, and atypical DUSPs make up the dual specificity phosphatases (DUSPs) superfamily (Patterson et al., 2009). DUSPs, characterized by their unique ability to dephosphorylate both tyrosine and serine/threonine residues within one substrate, are important modulators of multiple signaling pathways that regulate cellular processes like proliferation, apoptosis, and migration (Kim et al., 2016; Patterson et al., 2009). As a known cytoplasmic enzyme that exhibits a preferential selectivity towards ERK1/2 (Dowd et al., 1998; Keyse, 2008; Owens and Keyse, 2007), DUSP7 has been reported to be involved in the ERK signaling pathway and is an essential regulator of oocyte meiosis (Caunt et al., 2008; Pfender et al., 2015; Tischer and Schuh, 2016). DUSP7 contains a Rhodanese-like domain at its N terminus and a dual phosphatase domain at its C terminus. Two key amino acid residues within the conserved catalytic sequence (H/V)C(*X*5)R(S/T) of the phosphatase domain, C331 and R337, are important for DUSP7’s phosphatase activity (Denu and Dixon, 1995; Zhang et al., 1994). However, in contrast to other MKPs like DUSP6 and DUSP9, little is known about the physiological functions of DUSP7.

To analyze the function of DUSP7 in cell division, we generated the DUSP7 protein association and proximity networks and identified ERK2 as a mitotic interactor. Binding experiments showed that DUSP7 bound to ERK2 independently of ERK2 phosphorylation. Overexpression of DUSP7, but not catalytic inactive DUSP7, led to a decrease in phospho-ERK2 levels and chromosome alignment defects. DUSP7 depleted cells also displayed abnormalities in chromosome alignment and segregation and a slowed progression through mitosis. Chemical inhibition of MEK (leading to a decrease in active phospho-ERK2) or ERK2 (leading to a decrease in ERK2 kinase activity) led to errors in chromosome alignment. These data support a model where MEK phosphorylation activity and DUSP7 phosphatase activity regulate the levels of active phospho-ERK2, which is important for the fidelity of chromosome alignment and segregation during cell division.

## RESULTS

### DUSP7 interacts with ERK2 and regulates the levels of phospho-ERK2

During interphase, DUSP7 is an important factor in the ERK signaling pathway and exhibits a preferential selectivity towards ERK2 dephosphorylation (Dowd et al., 1998; Keyse, 2008; Owens and Keyse, 2007). To better understand the role of DUSP7 during cell division, we began by defining the protein-protein interaction network and protein proximity network of DUSP7 in mitotic cells. Localization and affinity purification (LAP= GFP-Tev-S-tag)-tagged and biotin identification 2 (BioID2)-tagged DUSP7 inducible HeLa stable cell lines were used to express LAP/BioID2-DUSP7 and biochemical purifications were analyzed by mass spectrometry (see STAR Methods for details). In-house R scripts were used to analyze the mass spectrometry data; the resultant protein interaction and proximity networks were visualized with RCytoscape JS (see STAR Methods for details) (Figures S1A – B). Further, we applied Gene Ontology (GO) terms (mitotic spindle; kinetochore and chromosome segregation) and CORUM complex annotation analyses to these networks (Figures S1C, 1A – B) (see STAR Methods for details). These analyses determined that ERK2 (aka MAPK1) was also associating with DUSP7 in mitosis (Figures 1A – B). Next, we validated the DUSP7-ERK2 mitotic interaction by immunoprecipitation (IP) experiments using mitotic cell extracts from Taxol-or Nocodazole-arrested LAP-DUSP7 stable cell lines (Figure 1C).

**Figure 1.**
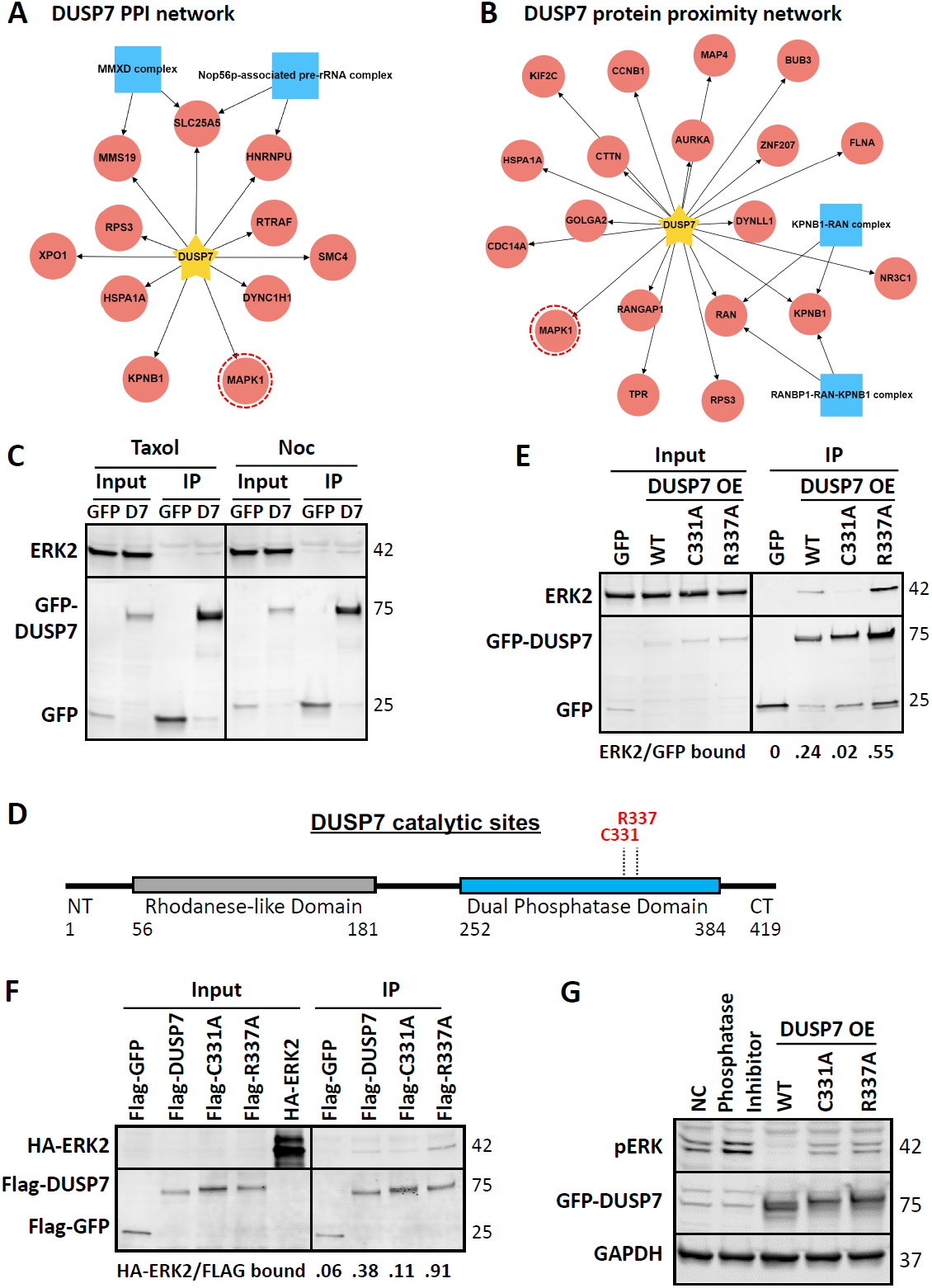
DUSP7 interacts with ERK2 and regulates the levels of phospho-ERK2. (A – B) DUSP7 protein-protein interaction (PPI) network (A) and protein proximity network (B) generated using mitotic spindle GO annotations and CORUM complex annotation analyses. Yellow star indicates the bait protein DUSP7; red circles indicate the potential interactors; blue squares indicate protein complex; red dashed circle highlights ERK2. (C) DUSP7 interacts with ERK2 in mitosis. LAP-only and LAP-DUSP7 HeLa stable cell lines were induced with 0.1μg/ml Doxycycline and arrested in mitosis by 100nM Taxol or 330nM Nocodazole for 18 hours before being harvested for S-tag pull downs. Pull downs were resolved by SDS PAGE, transferred to a PVDF membrane, and immunoblotted with indicated antibodies. (D)Schematic of DUSP7 domain structure and catalytic sites. Grey box represents Rhodanese-like domain, blue box represents dual phosphatase domain. NT= N terminus, CT= C terminus. The numbers of amino acid residues are indicated for each domain. DUSP7 catalytic sites are highlighted in red. (E – F) DUSP7-ERK2 interaction is influenced by DUSP7’s catalytic activity. In (E) LAP-only, LAP-DUSP7-WT, LAP-C331A, and LAP-R337A HeLa stable cell lines were induced with 0.1μ g/ml Doxycycline for 18 hours before being harvested for S-tag pull downs. In (F) HA-ERK2, Flag-DUSP7, Flag-C331A, Flag-R337A and Flag-GFP (negative control) were expressed in an IVT (*In Vitro* Transcription/Translation) system and incubated with anti-FLAG M2 magnetic beads in immunoprecipitation assays. Pull downs or IPs were resolved by SDS PAGE, transferred to a PVDF membrane, and immunoblotted with indicated antibodies. Ratios below the immunoblots indicate the relative protein-protein binding affinity. (G) DUSP7 regulates the levels of phospho-ERK2. HeLa cells were transfected with DUSP7, C331A, or R337A cDNA before being lysed and analyzed by immunoblot. Phosphatase inhibitor in the second lane was added when lysing the cells. Numbers on the right side of the immunoblots indicate the molecular weight of the proteins. OE= overexpression, WT= wild type, NC= negative control.

To better understand the DUSP7-ERK2 interaction, we generated a series of LAP-DUSP7 stable cell lines expressing either DUSP7 truncations or the C331A or R337A catalytic dead mutants (Figures S1F and 1D). IP experiments using cell extracts from these cell lines showed that ERK2 IPed with wild type DUSP7 and DUSP7-R337A but not DUSP7-C331A (Figure 1E). Similar results were obtained through *in vitro* binding experiments (Figure 1F). Furthermore, ERK2 failed to associate with DUSP7 truncations (Figure S1G), suggesting that full-length DUSP7 was necessary for ERK2 binding. Next, we sought to determine the significance of the DUSP7-ERK2 interaction. While overexpression of DUSP7 led to the absence of phospho-ERK2, overexpressed DUSP7-R337A or DUSP7-C331A showed a reduced ability to dephosphorylate ERK2 in HeLa cells (Figure 1G). Together, these results showed that DUSP7 was binding to ERK2 during mitosis and that DUSP7 was regulating the levels of active phospho-ERK2 through its phosphatase activity.

### Knockdown of DUSP7 leads to chromosome alignment and segregation defects

To understand the importance of DUSP7’s function in regulating the levels of active phospho-ERK2 during cell division, we first identified siRNA’s capable of depleting DUSP7 protein levels by immunoblot analysis and DUSP7 mRNA expression by RT-qPCR (Figures 2A and S2A – C). We then analyzed the consequences of depleting DUSP7 during cell division with fixed-cell immunofluorescence (IF) microscopy (Figures 2B and 2E). This analysis showed that depletion of DUSP7 led to an increased percentage of defective mitotic cells with chromosome misalignment (siDUSP7= 44.6±5.6, p<.05 compared to siControl= 29.1±2.9) (Figures 2C – D). DUSP7 depletion also led to defects in spindle organization including unfocused and multipolar spindles (Figure 2C). The chromosome misalignments defects in siDUSP7 cells translated into an approximate two-fold increase in the percentage of lagging chromosomes during anaphase (siDUSP7= 24.5±5.0, compared to siControl= 13.5±5.1) (Figure 2F). Next, we analyzed whether depletion of DUSP7, and its associated defects in chromosome alignment, was affecting the timing of cell division by live-cell time-lapse microscopy in HCT116 expressing GFP-H2B cells (Figure 2G). This analysis showed that depletion of DUSP7 led to marked increase in the time from chromosome condensation to chromosome segregation (siDUSP7= 54.0±38.3 minutes, p<.01 compared to siControl= 38.0±19.1 minutes) (Figures 2H – J; Movies S1 – S4). Together, these results showed that depletion of DUSP7 led to a slowed mitosis where cells failed to properly align and segregate chromosomes.

**Figure 2.**
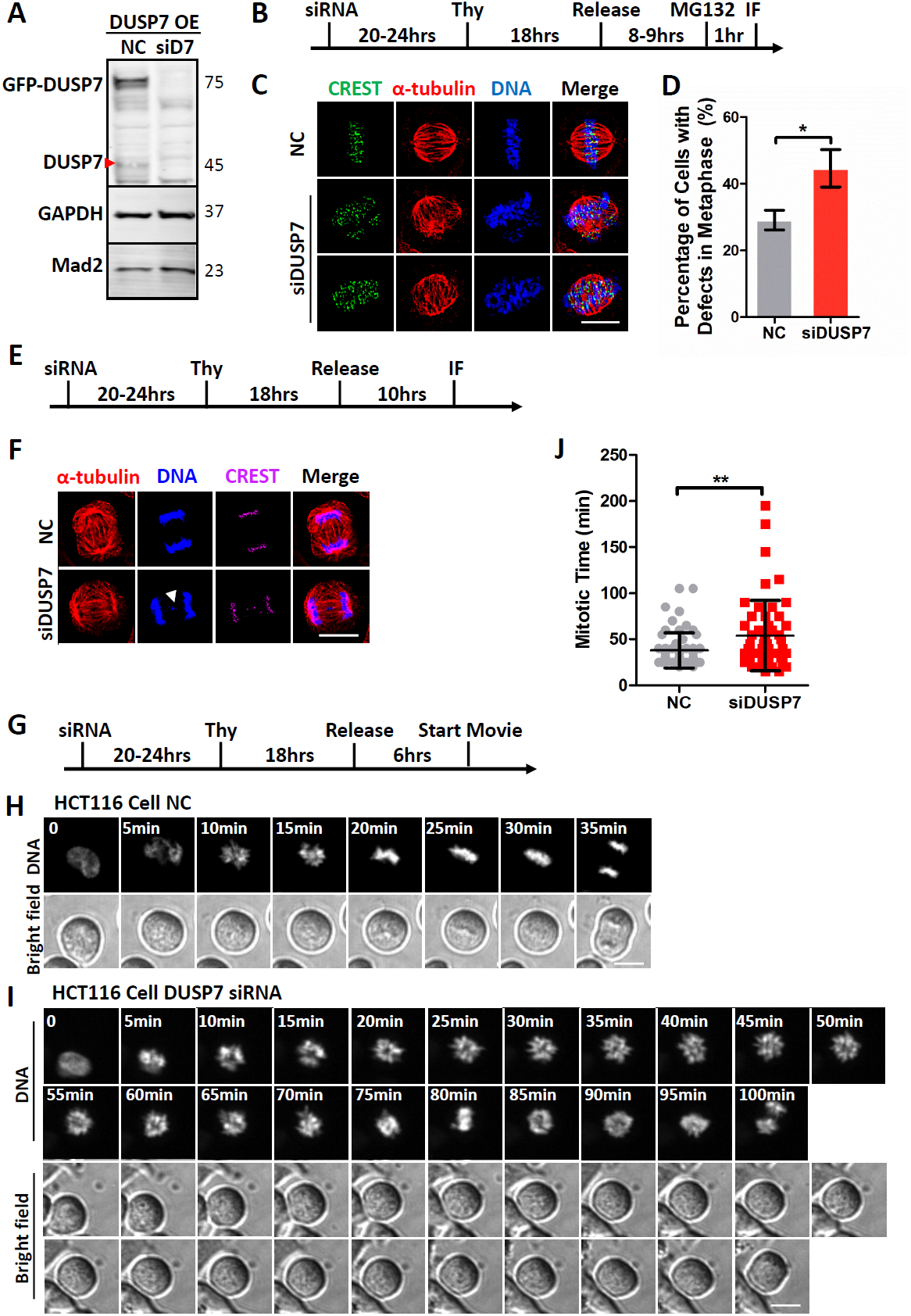
Knockdown of DUSP7 leads to chromosome alignment and segregation defects. (A) siRNA knockdown of DUSP7. LAP-DUSP7 HeLa stable cell lines were transfected with control (NC) or DUSP7 siRNA (siD7) for 72 hours before being lysed and analyzed by immunoblot. Numbers on the right side of the immunoblots indicate the molecular weight of the proteins. Red arrow indicates endogenous DUSP7 band. (B) Schematic of the immunofluorescence microscopy experiment performed in (C). (C) Knockdown of DUSP7 leads to chromosome misalignment in metaphase. HeLa cells were treated with negative control siRNA or siDUSP7 before being fixed and co-stained with anti-CREST and anti-α-tubulin antibodies and the DNA dye Hoechst 33342. Scale bar: 10μm. (D) Quantification of the percentage of cells with chromosome misalignment in metaphase (y-axis) for the conditions shown in (C) (x-axis). Three independent experiments were performed with about 300 cells in total counted for each quantification. Data are represented as mean±SD and * indicates P< 0.05. (E) Schematic of the immunofluorescence microscopy experiment performed in (F). (F) Knockdown of DUSP7 leads to an increase in lagging chromosomes in anaphase. HeLa cells were treated with negative control siRNA or siDUSP7 before being fixed and co-stained with anti-CREST and anti-α-tubulin antibodies and the DNA dye Hoechst 33342. Scale bar: 10μm. White arrow shows the lagging chromosome. (G) Schematic of the live-cell time-lapse microscopy experiment performed in (H) and (I). (H – I) Knockdown of DUSP7 leads to a prolonged time of chromosome congression during mitosis. Live-cell time-lapse microscopy of HCT116 cells treated with negative control siRNA (H) and siDUSP7 (I) undergoing cell division. Scale bars: 10μm. (J) Quantification of the timing of mitosis from chromosome condensation to chromosome segregation (y-axis) for the conditions shown in (H) and (I) (x-axis). Three independent experiments were performed with about 100 cells in total counted for each quantification. Data are represented as mean±SD and ** indicates P< 0.01. OE= overexpression, NC= negative control, siD7= siDUSP7, Thy= thymidine.

### DUSP7 promotes chromosome alignment in mitosis by regulating the activity of ERK2

ERK2 has an established role as a regulator of the G1/S transition, but whether or not ERK2 plays an important role in mitosis has been controversial (Meloche and Pouyssegur, 2007). Therefore, we sought to determine if ERK2 was important for human cell division. HeLa cells were treated with the MEK selective inhibitor U0126 (Duncia et al., 1998; Favata et al., 1998) or the ERK2 ATP-competitive inhibitor FR180204 (Ohori et al., 2005) and analyzed by IF microscopy (Figure 3A). In comparison to the control DMSO treatment, cells treated with U0126 or FR180204 showed a significant increase in chromosome alignment errors (U0126= 48.7±12.7, p<.05 and FR 180204= 45.6±6.5, p<.05 compared to DMSO= 25.8±3.9) (Figures 3B – C). These results showed that inhibiting ERK2 phosphorylation, and thereby its activation, or ERK2’s kinase activity led to severe chromosome alignment defects.

**Figure 3.**
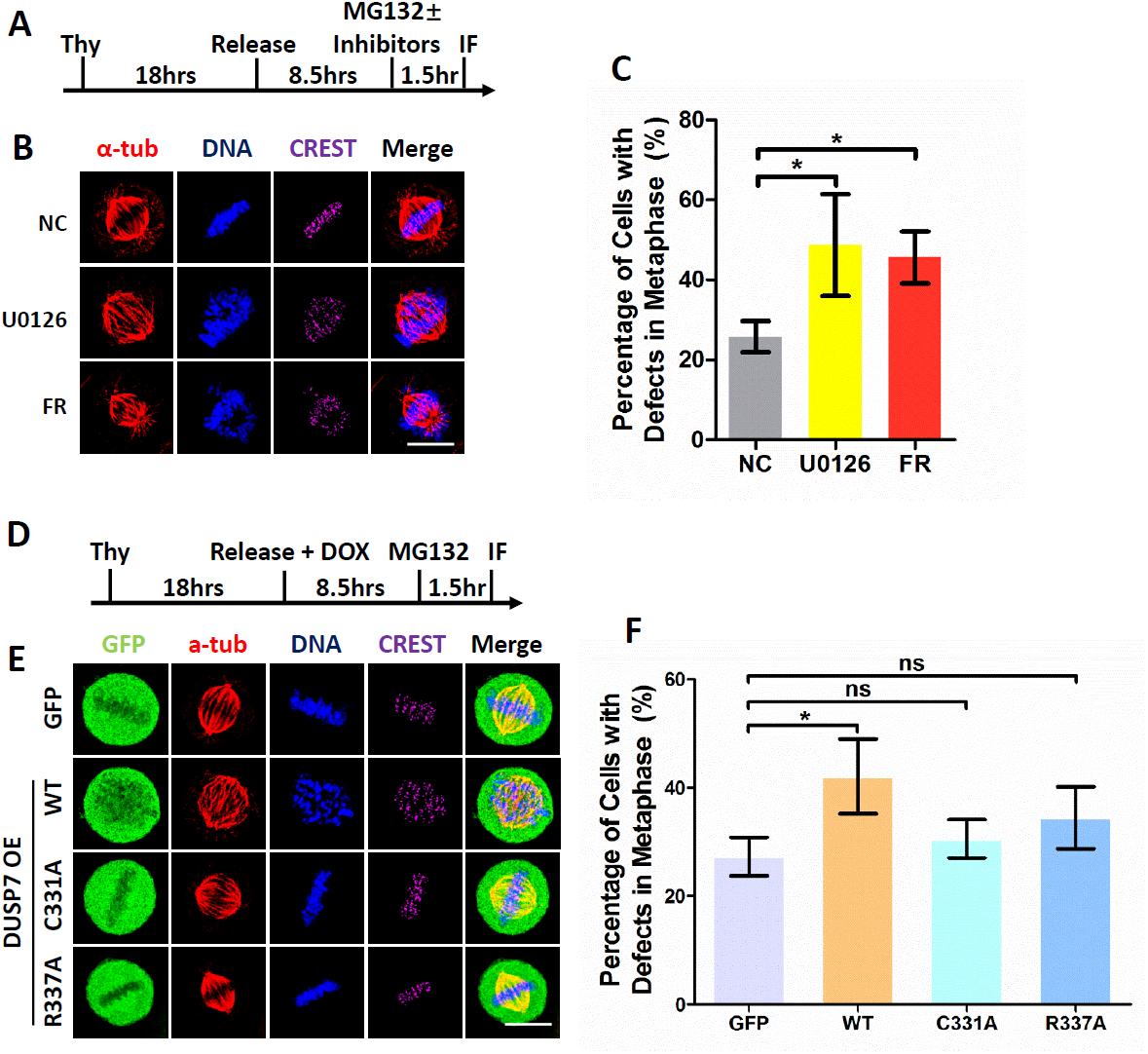
DUSP7 promotes chromosome alignment in mitosis by regulating ERK2 activity. (A) Schematic of the immunofluorescence microscopy experiment performed in (B). (B) Inhibition of MEK kinase activity (with 50μM U0126) or ERK2 kinase activity (with 50μM FR 180204) leads to chromosome misalignment in metaphase. HeLa cells were treated with DMSO (NC) or the indicated inhibitors, fixed, and co-stained with anti-CREST and anti-α-tubulin antibodies and the DNA dye Hoechst 33342. Scale bar: 10μm. (C) Quantification of the percentage of cells with chromosome misalignment in metaphase (y-axis) for the conditions shown in (B) (x-axis). Three independent experiments were performed with about 300 cells in total counted for each quantification. Data are represented as mean±SD and * indicates P< 0.05. (D) Schematic of the immunofluorescence microscopy experiment performed in (E). (E) Overexpression of DUSP7 wild type, but not catalytic dead mutants, leads to chromosome misalignment in metaphase. LAP-only, LAP-DUSP7-WT, LAP-C331A and LAP-R337A HeLa stable cell lines were treated as described in (D) before being fixed and co-stained with anti-GFP, anti-CREST, and anti-α-tubulin antibodies and the DNA dye Hoechst 33342. Scale bar: 10μm. (F) Quantification of the percentage of cells with chromosome misalignment in metaphase (y-axis) for the conditions shown in (E) (x-axis). Three independent experiments were performed with about 300 cells in total counted for each quantification. Data are represented as mean±SD and * indicates P< 0.05, ns indicates not statistically significant. Thy= thymidine, NC= negative control, FR= FR 180204.

Since DUSP7 shows a selective preference towards ERK2 and dephosphorylates ERK2 (Figure 1G), we hypothesized that overexpression of DUSP7 would lead to similar chromosome alignment defects to those observed in cells treated with the MEK inhibitor U0126. To test this, we overexpressed GFP-tagged DUSP7 (validated to decrease phospho-ERK2 levels, Figure 1G) or the catalytic dead DUSP7-C331A or DUSP7-R337A mutants (showed minimal effects on phospho-ERK2 levels, Figure 1G) and analyzed the cells by IF microscopy (Figure 3D). While DUSP7 overexpression led to a significant increase in chromosome alignment defects, overexpression of the DUSP7-R337A or DUSP7-C331A mutants did not (DUSP7= 42.1±6.9, p<.05; DUSP7-C331A= 30.6±3.5, p<.5; and DUSP7-R337A= 34.4±5.7, p<.5; compared to GFP control= 27.3±3.5) (Figures 3E – F). These results showed that an overabundance of DUSP7 phosphatase activity led to chromosome alignment defects.

### ERK2 interacts with DUSP7 independently of ERK2 phosphorylation

To further understand the ERK2-DUSP7 interaction, we sought to determine if it was dependent on ERK2 phosphorylation. IP experiments using cell extracts from a U0126-treated LAP-DUSP7 stable cell line showed that ERK2 bound to DUSP7 in the absence of MEK kinase activity (Figure 4A). Since ERK2 is phosphorylated by MEK at T185 and Y187 (Haystead et al., 1992; Payne et al., 1991), we generated the non-phosphorylation mimetic mutant ERK2-2A (T185A/Y187A) and the phosphorylation mimetic mutants ERK2-2D (T185D/Y187D) and ERK2-2E (T185E/Y187E) and analyzed their binding to DUSP7. *In vitro* binding experiments showed that ERK2, ERK2-2A, ERK2-2D and ERK2-2E all bound to DUSP7 with no major changes in binding (Figure 4B). Similar results were observed in IP experiments from HeLa cell extracts (Figure 4C). Together, these results showed that DUSP7’s binding to ERK2 did not require ERK2 to be phosphorylated.

**Figure 4.**
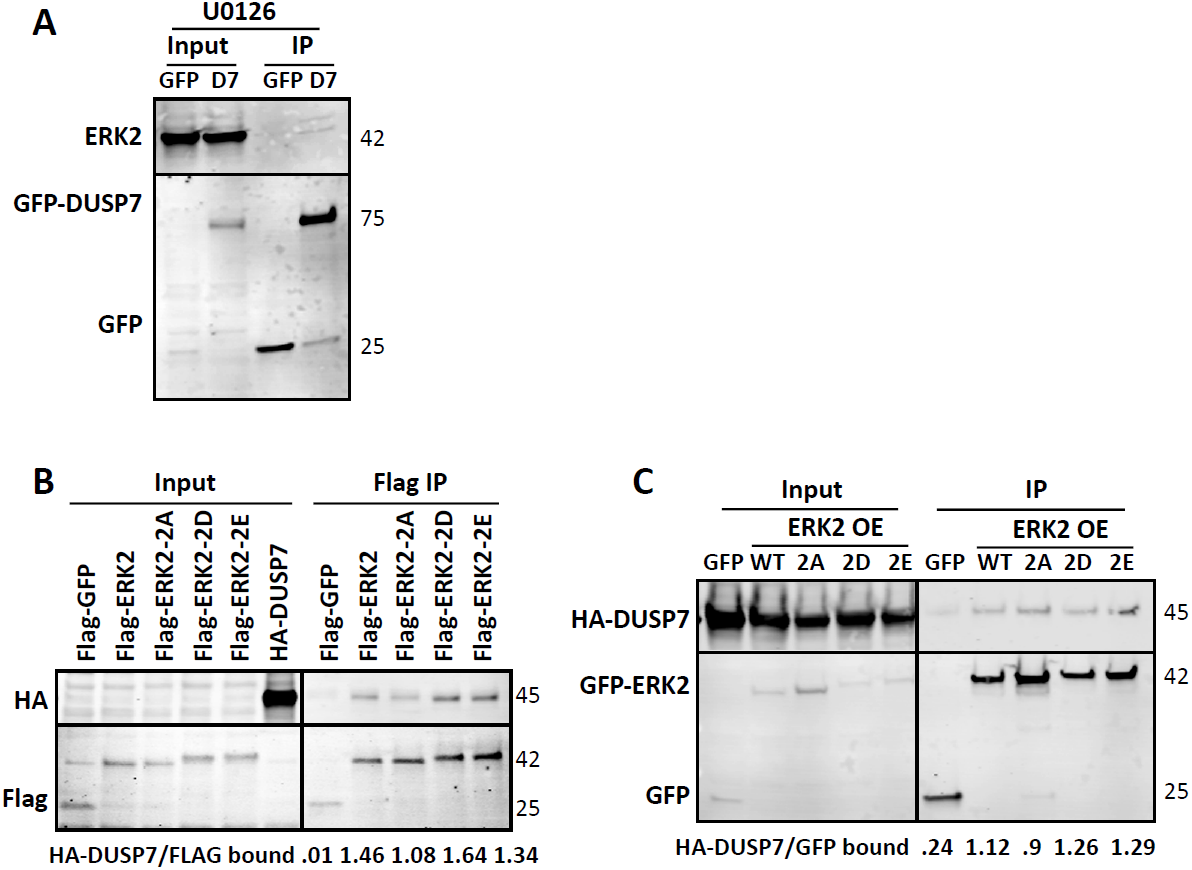
ERK2 interacts with DUSP7 independently of ERK2 phosphorylation. (A) LAP-only and LAP-DUSP7-WT HeLa stable cell lines were induced by 0.1μg/ml Doxycycline and inhibited by 50μM U0126 for 18 hours before being harvested for S-tag pull downs. Pull downs were resolved by SDS PAGE, transferred to a PVDF membrane, and immunoblotted with the indicated antibodies. (B) HA-DUSP7, Flag-ERK2, Flag-ERK2-2A, Flag-ERK2-2D, Flag-ERK2-2E and Flag-GFP (as the negative control) were expressed in an IVT (*In Vitro* Transcription/Translation) system and incubated with anti-FLAG M2 magnetic beads in immunoprecipitation assays. IPs were resolved by SDS PAGE, transferred to a PVDF membrane, and immunoblotted with indicated antibodies. (C) LAP-only, LAP-ERK2-WT, LAP-ERK2-2A, LAP-ERK2-2D and LAP-ERK2-2E HeLa stable cell lines were transiently transfected with HA-DUSP7 and induced by 0.1μg/ml (A) Doxycycline for 18 hours before being harvested for S-tag beads pull down. Pull downs were resolved by SDS PAGE, transferred to a PVDF membrane, and immunoblotted with indicated antibodies. Numbers on the right side of the immunoblots indicate the molecular weight of the proteins. Ratios below the immunoblots (B) and (C) indicate the relative protein-protein binding affinity. OE= overexpression, Noc= Nocodazole, D7= DUSP7, 2A= T185A/Y187A, 2D= T185D/Y187D, 2E= T185E/Y187E, WT= wild type.

## DISCUSSION

This study aimed to advance our understanding of DUSP7’s function. Our data are consistent with a model where during a normal mitosis, the MEK kinase activity phosphorylates ERK2 and DUSP7’s phosphatase activity dephosphorylates ERK2 to establish an equilibrium of active phospho-ERK2. This phospho-ERK2 equilibrium is critical for ensuring the fidelity of chromosome alignment and segregation. Consistent with this model, mass spectrometry analyses of DUSP7 LAP-based affinity purifications and DUSP7 BioID2-based proximity labeling purifications showed that DUSP7 was associating with ERK2 during mitosis. This result was further confirmed by in cell and *in vitro* binding experiments. Importantly, pharmacological inhibition of MEK (leading to a decrease in active phospho-ERK2) or inhibition of ERK2 (leading to a decrease in ERK2 kinase activity) led to errors in chromosome alignment. Additionally, overexpression of DUSP7, but not DUSP7 catalytic dead mutants, led to a decrease in phospho-ERK2 and chromosome alignment defects. Furthermore, depletion of DUSP7 by RNAi also led to abnormalities in chromosome alignment and segregation and to a slowed progression from chromosome condensation to chromosome segregation. Together, these results demonstrate that the MAPK/ERK pathway is important for cell division and that DUSP7 is an important regulator of this pathway. These studies establish DUSP7 as an important mitotic phosphatase that regulates the abundance of active phospho-ERK2 to ensure the fidelity of chromosome alignment and segregation.

Of interest, we found that ERK2 and DUSP7 both localized throughout the cell during cell division with weak spindle localization in prometaphase and metaphase (Figure S3A – C). However, the subcellular localization of phospho-ERK has remained controversial with studies reporting it as kinetochore and centrosome localized, which can be diminished by inhibiting the phospho-ERK antibody with the phosphopeptide but not by inhibiting MEK activity with U0126 (Shapiro et al., 1998; Shinohara et al., 2006; Zecevic et al., 1998). Using five different commercial anti-phospho-ERK antibodies, we observed different phospho-ERK subcellular localizations (Figure S3F – J). Except for the antibody from CST (Cat# 4377) (Figure S3H), the other four anti-phospho-ERK antibodies showed constant phospho-ERK subcellular localizations in the presence or absence of U0126 or FR 180204 treatments (Figure S3F, S3G, S3I and S3J), even though all antibodies were able to detect a decrease in phospho-ERK2 protein levels in MEK inhibited (U0126-treated) cells (Figure S3D – E). Therefore, caution should be taken when making inferences about the localization of phospho-ERK2 during cell division when using anti-phospho-ERK antibodies.

With the exception of ERK2, there is little known about the repertoire of DUSP7 substrates, regulators, and interactors. The GO enrichment analyses of DUSP7 protein association network and DUSP7 proximity protein network indicate that DUSP7 is likely to associate with numerous proteins that carry out important functions related to a broad array of cellular processes including apoptotic process, the regulation of transcription, and cell division (Figure S1D – E). Therefore, future studies aimed at understanding the importance of these interactions will further aid our understanding of DUSP7’s function in cell division and beyond.

We note that the dysregulation of DUSP protein levels has been linked to the initiation and development of various types of cancers (Bermudez et al., 2010; Keyse, 2008; Nunes-Xavier et al., 2011). More specifically, DUSP7 has been found to be overexpressed in blood cancers, including acute lymphoblastic leukemia (ALL) and acute myeloid leukemia (AML) (Levy-Nissenbaum et al., 2003a; Levy-Nissenbaum et al., 2003b, c). Of interest, we have also observed elevated DUSP7 protein levels in multiple blood cancer cell lines including ALL CCRF-CEM cells (Torres lab unpublished). Therefore future investigations related to DUSP7’s role in ensuring the fidelity of chromosome movements during cell division within the context of acute leukemia could advance our understanding of the development of blood cancers.

## STAR☆METHODS

Detailed methods are provided in the online version of this paper and include the following:

- **KEY RESOURCES TABLE**
- **LEAD CONTACT AND MATERIALS AVAILABILITY**
- **METHODS DETAILS**
  - Cell culture
  - Plasmids, mutagenesis, and generation of stable cell lines
  - LAP/BioID2 purifications and LC-MS/MS analyses
  - Immunoprecipitations, *in vitro* binding assays, and immunoblot analyses
  - Immunofluorescence and live-cell time-lapse microscopy
  - RT-qPCR
  - Antibodies
- **QUANTIFICATION AND STATISTICAL ANALYSIS**
- **DATA AND CODE AVAILABILITY**

## SUPPLEMENTAL INFORMATION

Supplemental Information includes Supplemental Experimental Procedures, 3 figures, and 4 movies.

## ACKNOWLEDGEMENTS

This material is based upon work supported by the National Institutes of Health NIGMS grant number R01GM117475 to J.Z.T., any opinions, findings, and conclusions or recommendations expressed in this material are those of the authors and do not necessarily reflect the views of the National Institutes of Health NIGMS. Y.A.G. was supported by the UCLA Tumor Cell Biology Training Program (USHHS Ruth L. Kirschstein Institutional National Research Service Award # T32CA009056). This work was supported in part by a grant to The University of California, Los Angeles from the Howard Hughes Medical Institute through the James H. Gilliam Fellowships for Advanced Study Program (E.F.V), by a UCLA Molecular Biology Institute Whitcome Fellowship (E.F.V.) and a NIH P30 DK063491 grant (J.P.W.). Work performed in the UCLA Molecular Screening Shared Resource was supported by the National Cancer Institute of the National Institutes of Health under award number P30CA016042.

## AUTHOR CONTRIBUTIONS

XG, YAG, IR, EV, LWG, AAG, EV, JPW, BT, RD and JZT performed experiments and discussed results. XG and JZT wrote the paper with input from YAG, IR, EV, LWG, AAG, EV, JPW, BT and RD.

## Graphic Abstract, Guo et al., 2020

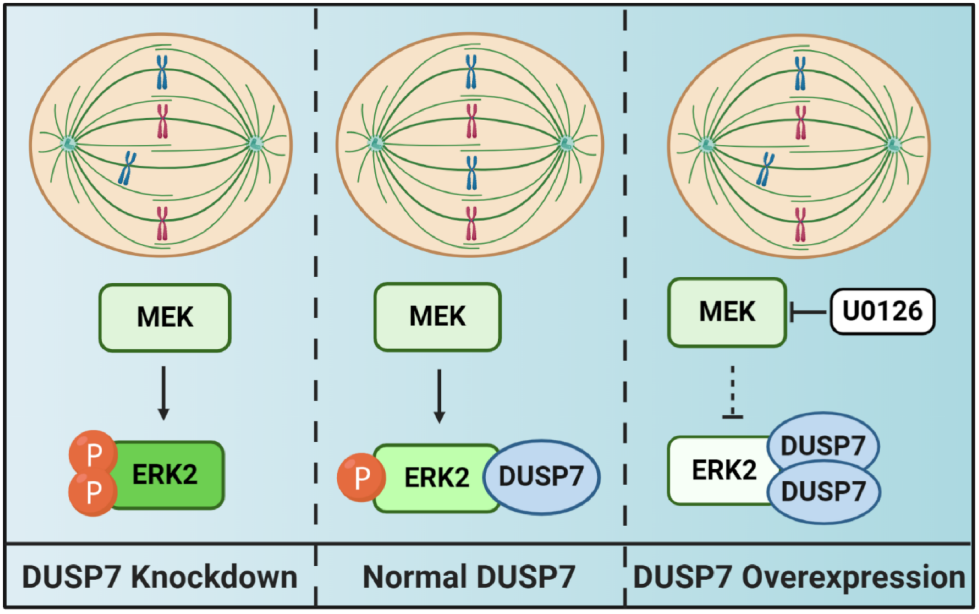

