## Supplemental Material for "DUSP7 Regulates the Activity of ERK2 to Promote Proper Chromosome Alignment During Cell Division"

#### SUPPLEMENTAL FIGURES

**Figure S1. Mass spectrometry analysis of DUSP7 mitotic interactors and validation of the DUSP7-ERK2 binding. Related to Figure 1.**

(A) DUSP7 protein-protein interaction (PPI) network comprised of proteins identified in LAP-DUSP7 tandem affinity purification by mass spectrometry.

(B) DUSP7 protein proximity network comprised of proteins identified in BioID2-DUSP7 biotin purifications by mass spectrometry.

(C) GeneOntology (GO) terms used for generating PPI and protein proximity networks in Figure 1A and B.

(D – E) GO enrichment analyses of the DUSP7 protein-protein interaction network (D) and the DUSP7 proximity association network (E). Graphs show the most represented GO terms identified in the GO enrichment analysis.

(F) Schematic of DUSP7 domain truncations. The numbers of amino acid residues are indicated for each domain truncation. Grey box represents Rhodanese-like domain, blue box represents dual phosphatase domain. FL= full length, NT= N terminus, CT= C terminus, NTRD= N terminus and Rhodanese-like domain, RD= Rhodanese-like domain, PD= phosphatase domain, PDCT= phosphatase domain and C terminus.

(G) ERK2 does not interact with DUSP7 truncations. LAP-only, LAP-DUSP7-WT and the indicated truncations were established in HeLa cells and their expression was induced with 0.1μg/ml Doxycycline for 18 hours. Cells were harvested and protein extracts were used for S-tag pull downs, which were resolved by SDS PAGE, transferred to a PVDF membrane, and immunoblotted with the indicated antibodies. Numbers on the right side of the western blots indicate the molecular weight of the proteins. OE= overexpression, WT= wild type.

**Figure S2. siRNA knockdown of DUSP7. Related to Figure 2.**

(A) HeLa cells were transfected with control (NC) or DUSP7 siRNA (siD7) for 72 hours before being lysed and analyzed by immunoblot. Numbers on the right side of the immunoblots indicate the molecular weight of the proteins. Red arrow indicates endogenous DUSP7 band.

(B) and (C) The relative gene expression of DUSP7 and Mad2 (as a negative control to show siRNA's specificity towards DUSP7) in HeLa cells (B) or LAP-DUSP7 cell lines (C) were normalized by GAPDH and analyzed with the Livak-Schmittgen method ($2^{-\Delta\Delta Cq}$). Data represent the average ±SD of three independent experiments; technical triplicates were quantified for each experiment; * indicates $p < 0.05$, ** indicates $p < 0.01$, *** indicates $p < 0.001$, ns indicates not statistically significant. NC= negative control.

**Figure S3. Subcellular localization of ERK2, DUSP7 and phospho-ERK during cell division. Related to Figures 1 – 4.**

(A) Subcellular localization of overexpressed ERK2 during cell division. HeLa cells were transiently transfected with GFP/HA/Flag-ERK2, fixed and co-stained with anti-GFP, anti-HA, and anti-Flag antibodies and the DNA dye Hoechst 33342. Scale bars: 10µm.

(B) Subcellular localization of endogenous DUSP7 during cell division. HeLa cells were fixed and co-stained with anti-DUSP7, anti-α-tubulin, and anti-CREST antibodies and the DNA dye Hoechst 33342. Scale bar: 10µm.

(C) Subcellular localization of overexpressed DUSP7 during cell division. The LAP-DUSP7 HeLa stable cell line was induced with 0.1μg/ml Doxycycline for 18 hours before being fixed and co-stained with anti-GFP, anti-α-tubulin, and anti-CREST antibodies and the DNA dye Hoechst 33342. Scale bar: 10μm.

(D) Schematic of the western blot and immunofluorescence experiments performed in (E-J). Thy= thymidine, WB= western blot.

(E) Inhibition of MEK kinase activity but not ERK2 kinase activity reduces phospho-ERK levels. HeLa cells were treated as described in (D) before being harvested and lysed. The samples were resolved by SDS PAGE, transferred to a PVDF membrane, and immunoblotted with the indicated antibodies. Numbers on the right side of the western blots indicate the molecular weight of the proteins.

(F – J) Subcellular localization of phospho-ERK during cell division. HeLa cells were treated as described in (D) before being fixed and co-stained with anti-pERK (Santa Cruz (F), CST (4370S) (G), CST (4377) (H), Abcam (I), R&D (J)), and anti-CREST antibodies and the DNA dye Hoechst 33342. Scale bars: 10μm.

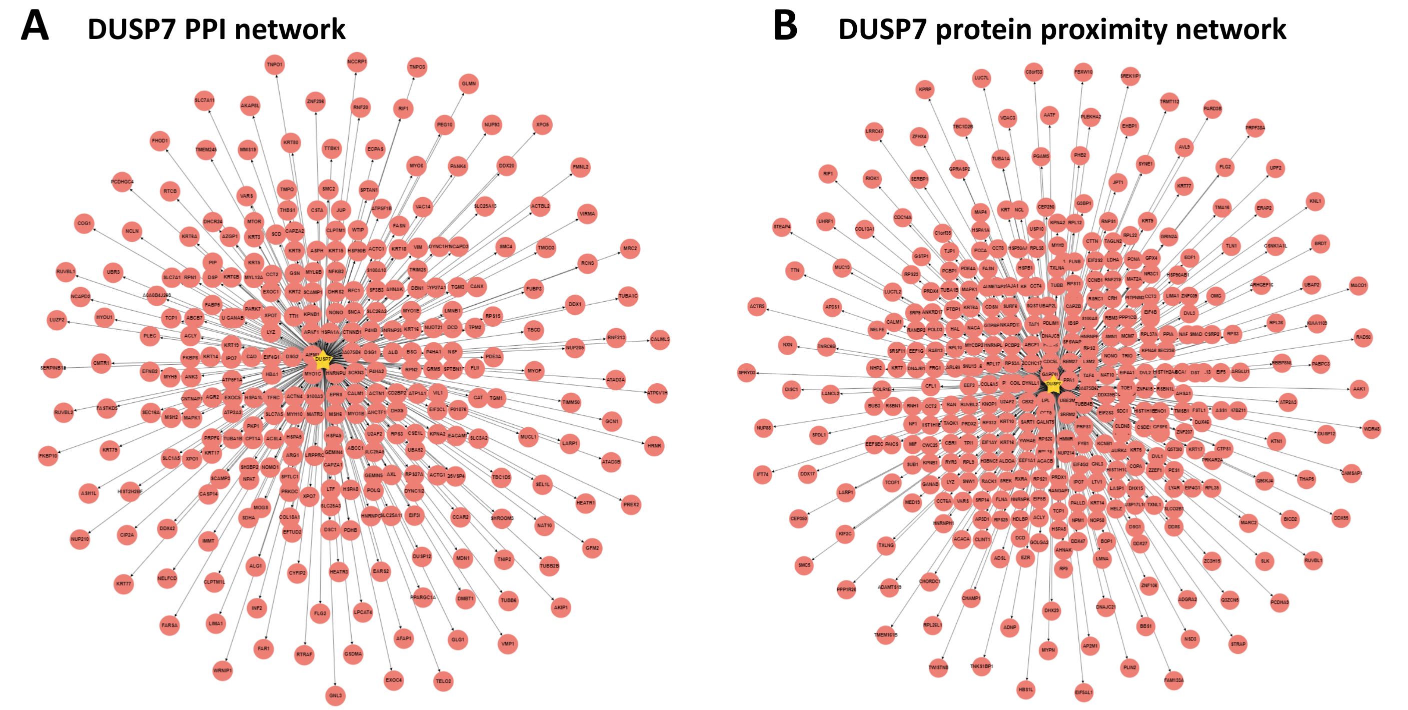


C

| GO:ID | Ontology |
| --- | --- |
| GO:0000776 | kinetochore |
| GO:0007079 | mitotic chromosome movement towards spindle pole |
| GO:0007052 | mitotic spindle organization |
| GO:1990023 | mitotic spindle midzone |
| GO:0090307 | mitotic spindle assembly |
| GO:0072686 | mitotic spindle |
| GO:1901673 | regulation of mitotic spindle assembly |
| GO:0040001 | establishment of mitotic spindle localization |
| GO:0000070 | mitotic sister chromatid segregation |
| GO:0007059 | chromosome segregation |

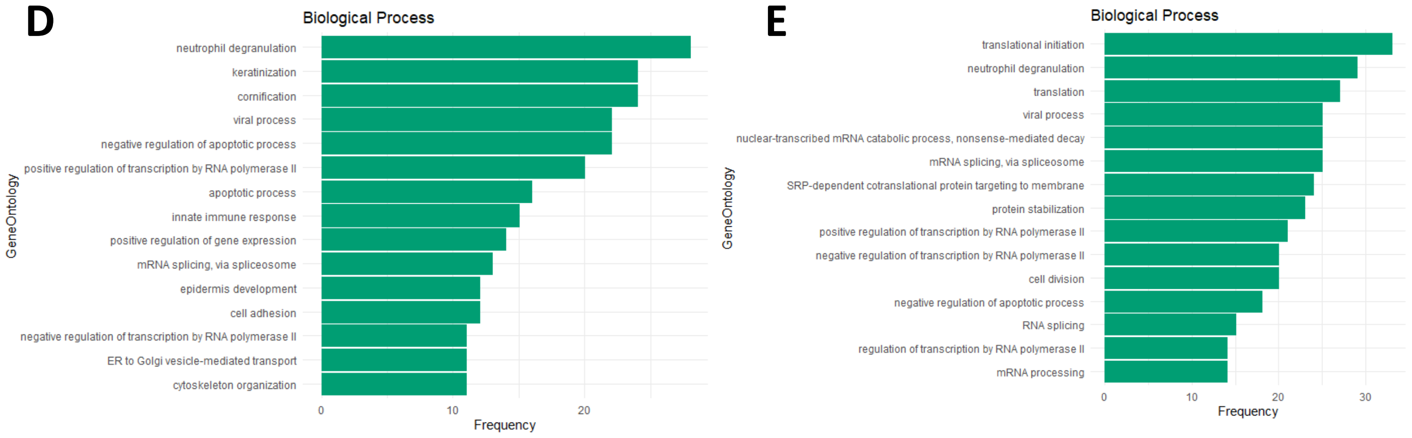


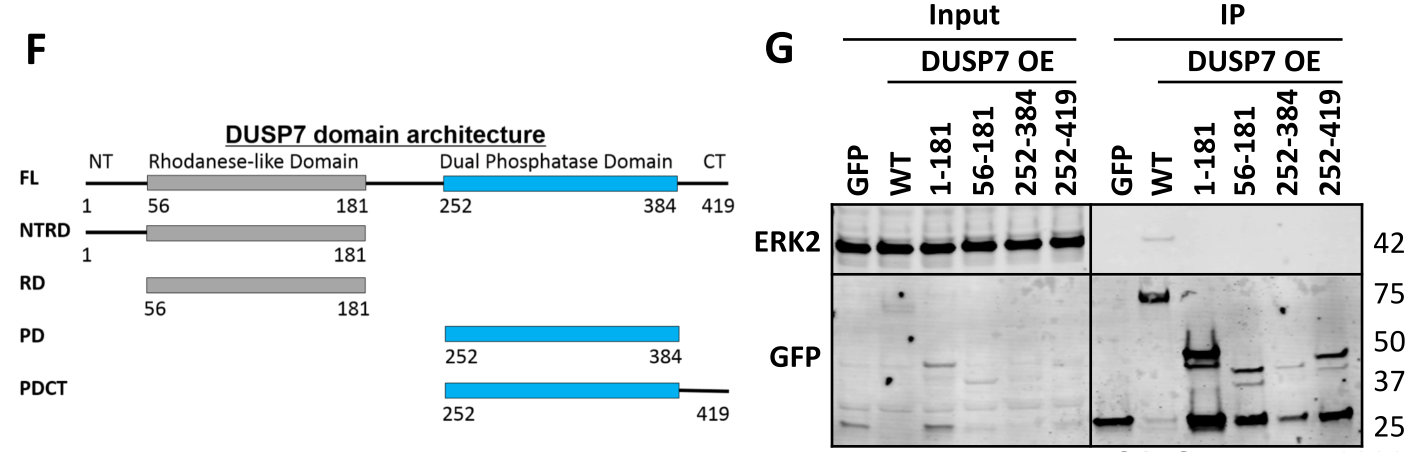


Figure S1, Guo et al., 2020

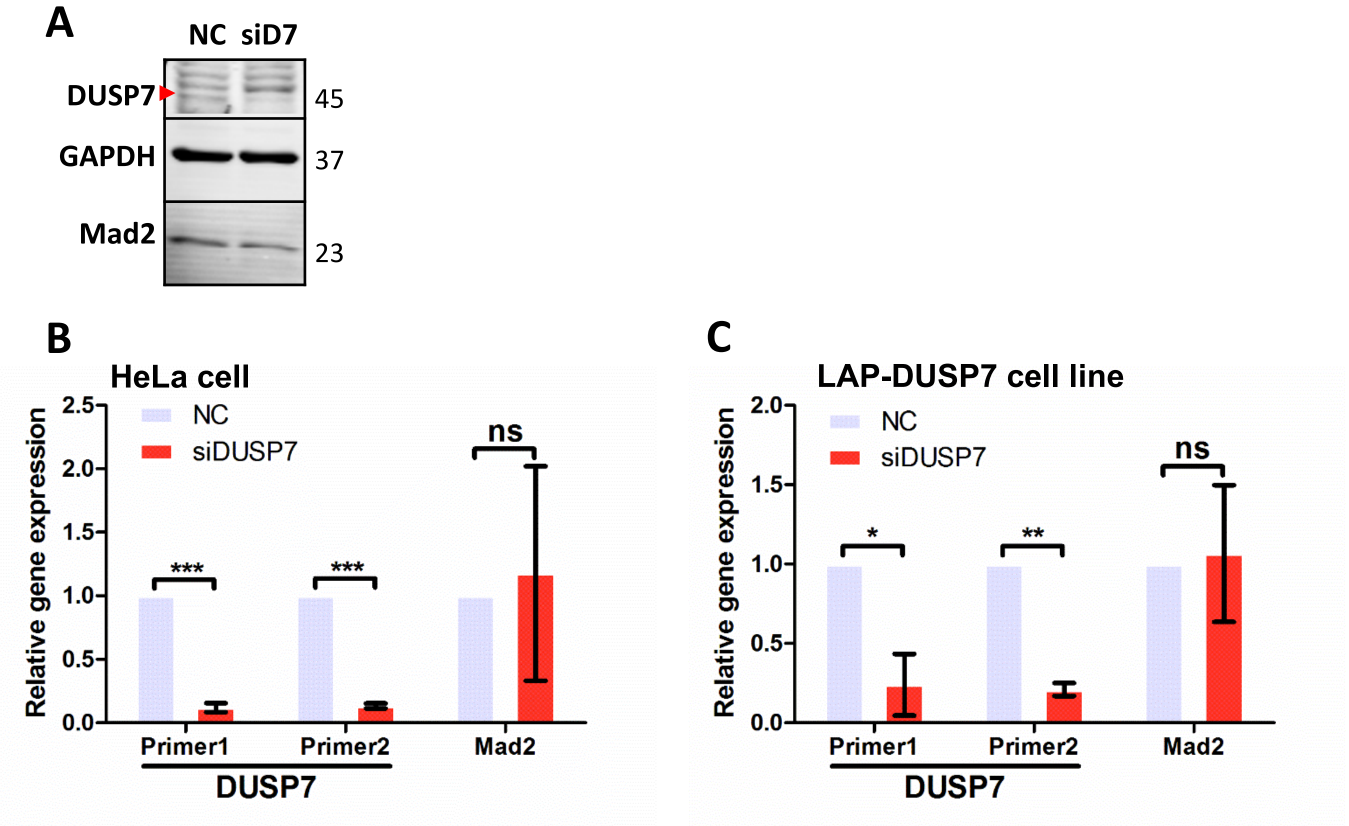


Figure S2, Guo et al., 2020

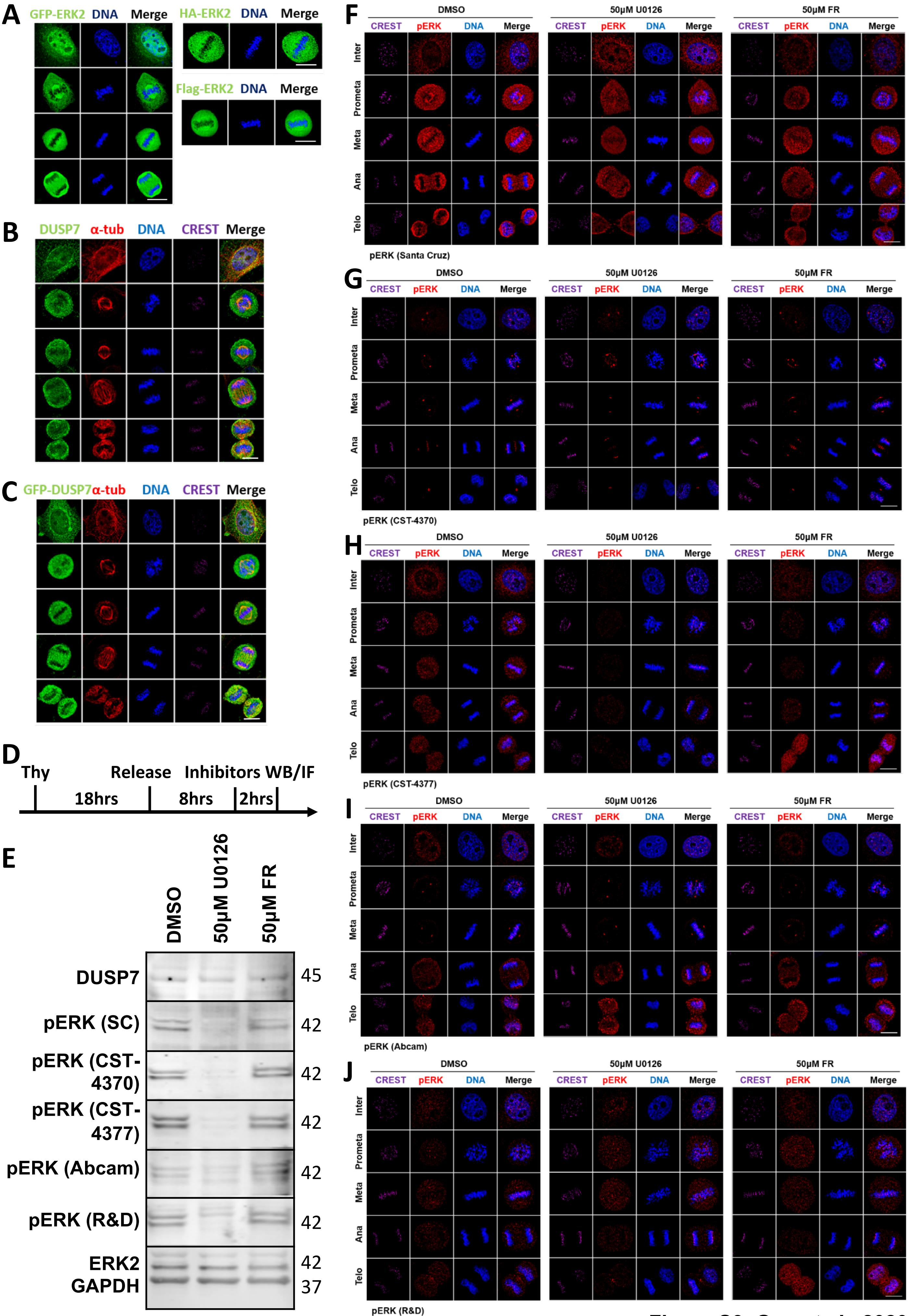


Figure S3, Guo et al., 2020

#### SUPPLEMENTAL MOVIES

**Movie S1 – 2. Live-cell time-lapse microscopy movie of a representative control HCT116 GFP-H2B cell undergoing cell division. Related to Figure 2.**

HCT116 GFP-H2B cells were mock transfected, treated as described in Figure 2H and imaged live six hours post thymidine release for 18 hours with an ImageXpress XL imaging system at 37 °C in 5% $CO_2$ using a 20x air objective. Images were captured every 5 minutes with bright field (Movie S1) and GFP (Movie S2) channels and converted to AVI movies with ImageJ at one frame per second. Each frame represents a five-minute interval.

**Movie S3 – 4. Live-cell time-lapse microscopy movie of representative siDUSP7 treated HCT116 GFP-H2B cell undergoing cell division. Related to Figure 2.**

HCT116 GFP-H2B cells were transfected with DUSP7 siRNA, treated as described in Figure 2H, and imaged live six hours post thymidine release for 18 hours with an ImageXpress XL imaging system at 37 °C in 5% $CO_2$ using a 20x air objective. Images were captured every five minutes with bright field (Movie S3) and GFP (Movie S4) channels and converted to AVI movies with ImageJ at one frame per second. Each frame represents a five-minute interval.

### STAR★METHODS

#### KEY RESOURCES TABLE

| REAGENT or RESOURCE | SOURCE | IDENTIFIER |
|---|---|---|
| Antibodies | | |
| Mouse monoclonal anti-ERK2 (D-2) | Santa Cruz Biotechnology | Cat# sc-1647; RRID: AB_627547 |
| Chicken polyclonal anti-GFP | Abcam | Cat# ab13970; RRID: AB_300798 |
| Rabbit polyclonal anti-HA | Proteintech | Cat# 51064-2-AP; RRID: AB_11042321 |
| Mouse monoclonal anti-FLAG® M2 | Sigma-Aldrich | Cat# F1804; RRID: AB_262044 |
| Mouse monoclonal anti-FLAG Dylight 800 Conjugated (clone 29E4.G7) | Rockland | Cat# 200-345-383; RRID: AB_10702994 |
| Rabbit monoclonal anti-phospho-p44/42 MAPK (Erk1/2) (Thr202/Tyr204) (clone D13.14.4E) | Cell Signaling Technology | Cat# 4370; RRID: AB_2315112 |
| Rabbit monoclonal anti-phospho-p44/42 MAPK (Erk1/2) (Thr202/Tyr204) (197G2) | Cell Signaling Technology | Cat# 4377; RRID: AB_331775 |
| Rabbit polyclonal anti-phospho-ERK1 (T202/Y204)/ERK2 (T185/Y187) | R&D Systems | Cat# AF1018; RRID: AB_354539 |
| Mouse monoclonal anti-p-ERK (E-4) | Santa Cruz Biotechnology | Cat# sc-7383; RRID: AB_627545 |
| Mouse monoclonal anti-phospho-ERK1/ERK2 (Thr185/Tyr187) (clone MAPK-YT) | Abcam | Cat# ab50011; RRID: AB_1603684 |
| Mouse monoclonal anti-GAPDH (clone 1E6D9) | Proteintech | Cat# 60004-1-Ig; RRID: AB_2107436 |
| Rabbit polyclonal anti-DUSP7/PYST2 | Proteintech | Cat# 26910-1-AP |
| Goat polyclonal anti-Mad2 (C-19) | Santa Cruz Biotechnology | Cat# sc-6329; RRID: AB_648599 |
| Human polyclonal anti-Centromere Protein/CREST | Antibodies Incorporated | Cat# 15-234-0001; RRID: AB_2687472 |
| Rat monoclonal anti-α-Tubulin (clone YOL1/34) | Bio-Rad | Cat# MCA78G; RRID: AB_325005 |
| Donkey polyclonal anti-Human IgG (H+L), Fluorescein (FITC) AffiniPure | Jackson ImmunoResearch Labs | Cat# 709-095-149; RRID: AB_2340514 |
| Donkey polyclonal anti-Rat IgG (H+L), Cy3 AffiniPure | Jackson ImmunoResearch Labs | Cat# 712-165-153; RRID: AB_2340667 |
| Donkey polyclonal anti-Human IgG (H+L), Cy5 AffiniPure | Jackson ImmunoResearch Labs | Cat# 709-175-149; RRID: AB_2340539 |
| Donkey polyclonal anti-Chicken IgY (IgG) (H+L), Fluorescein (FITC) AffiniPure | Jackson ImmunoResearch Labs | Cat# 703-095-155; RRID: AB_2340356 |
| Donkey polyclonal anti-Rabbit IgG (H+L), Cy3 AffiniPure | Jackson ImmunoResearch Labs | Cat# 711-165-152; RRID: AB_2307443 |
| Donkey polyclonal anti-Rabbit IgG (H+L), Fluorescein (FITC) AffiniPure | Jackson ImmunoResearch Labs | Cat# 711-095-152; RRID: AB_2315776 |

| | | |
|---|---|---|
| Donkey polyclonal anti-Mouse IgG (H+L), Fluorescein (FITC) AffiniPure | Jackson ImmunoResearch Labs | Cat# 715-095-151; RRID: AB_2335588 |
| Donkey polyclonal anti-Mouse IgG (H+L), Cy3 AffiniPure | Jackson ImmunoResearch Labs | Cat# 715-165-151; RRID: AB_2315777 |
| Donkey polyclonal anti-Goat IgG (H+L), IRDye 680RD | LI-COR Biosciences | Cat# 926-68074; RRID: AB_10956736 |
| Donkey polyclonal anti-Mouse IgG (H+L), IRDye 680RD | LI-COR Biosciences | Cat# 926-68072; RRID: AB_10953628 |
| Donkey polyclonal anti-Mouse IgG (H+L), IRDye 800CW | LI-COR Biosciences | Cat# 926-32212; RRID: AB_621847 |
| Donkey polyclonal anti-Chicken IgG (H+L), IRDye 800CW | LI-COR Biosciences | Cat# 926-32218; RRID: AB_1850023 |
| Donkey polyclonal anti-Rabbit IgG (H+L), IRDye 680RD | LI-COR Biosciences | Cat# 926-68073; RRID: AB_10954442 |
| Donkey polyclonal anti-Rabbit IgG (H+L), IRDye 800CW | LI-COR Biosciences | Cat# 926-32213; RRID: AB_621848 |
| Chemicals, Peptides, and Recombinant Proteins | | |
| U0126 | Selleckchem | Cat# S1102; CAS:1173097-76-1 |
| FR 180204 | Selleckchem | Cat# S7524; CAS:865362-74-9 |
| Paclitaxel | Sigma-Aldrich | Cat# T7191; CAS:33069-62-4 |
| Nocodazole | Sigma-Aldrich | Cat# 1404; CAS:31430-18-9 |
| Thymidine | Sigma-Aldrich | Cat# T1895; CAS:50-89-5 |
| Hygromycin B | Thermo Fisher Scientific | Cat# 10687010 |
| Doxycycline | Sigma-Aldrich | Cat# D9891; CAS:24390-14-5 |
| MG132 | Millipore Sigma | Cat# 474790; CAS:133407-82-6 |
| Halt Protease Inhibitor Cocktail | Thermo Fisher Scientific | Cat# 87786 |
| Hoechst 33342 | Thermo Fisher Scientific | Cat# H1399; CAS: 23491-52-3 |
| ProLong Gold Antifade Mountant | Thermo Fisher Scientific | Cat# P36934 |
| Lipofectamine RNAiMAX | Thermo Fisher Scientific | Cat# 13778150 |
| FuGENE HD Transfection Reagent | Promega | Cat# E2311 |
| FuGENE 6 Transfection Reagent | Promega | Cat# E2691 |
| S-protein Agarose | Millipore Sigma | Cat# 69704-4 |
| Anti-FLAG M2 magnetic beads | Sigma-Aldrich | Cat# M8823 |
| Biotin | Sigma-Aldrich | Cat# B4501-1G |
| Dynabeads MyOne Streptavidin C1 | Thermo Fisher Scientific | Cat# 65002 |
| Critical Commercial Assays | | |
| Gateway LR Clonase II Enzyme mix | Thermo Fisher Scientific | Cat# 11791020 |
| Gateway BP Clonase II Enzyme mix | Thermo Fisher Scientific | Cat# 11789020 |
| QuikChange Lightning Site-Directed Mutagenesis Kit | Agilent | Cat# 210518 |
| SP6 TnT Quick Coupled Transcription/Translation System | Promega | Cat# L2080 |
| PureYield Plasmid Miniprep System | Promega | Cat# A1222 |
| QIAprep Spin Miniprep Kit | QIAGEN | Cat# 27106 |
| PureYield Plasmid Midiprep System | Promega | Cat# A2495 |
| Direct-zol RNA Miniprep Kits | Zymo Research | Cat# R2051 |

| | | |
|---|---|---|
| UltraScript 2.0 cDNA Synthesis Kit | Genesee Scientific | Cat# 17-702 |
| qPCRBIO SyGreen Blue Mix Lo-ROX | Genesee Scientific | Cat# 17-505 |
| Deposited Data | | |
| Affinity-based mass spectrometry performed with LAP-DUSP7 | This paper | TBD |
| Proximity-based mass spectrometry performed with BioID2-DUSP7 | This paper | TBD |
| Experimental Models: Cell Lines | | |
| HeLa cells | ATCC | Cat# CCL-2; RRID: CVCL_0030 |
| HCT116 cells constitutively expressing GFP-H2B | This paper | N/A |
| HeLa Flp-In T-Rex cell lines | Stephen Taylor Lab | (Tighe et al., 2004) |
| Inducible HeLa LAP-DUSP7 stable cell line | This paper | N/A |
| Inducible HeLa BioID2-DUSP7 stable cell line | This paper | N/A |
| Inducible HeLa LAP-DUSP7-C331A stable cell line | This paper | N/A |
| Inducible HeLa LAP-DUSP7-R337A stable cell line | This paper | N/A |
| Inducible HeLa LAP-DUSP7-1-181aa stable cell line | This paper | N/A |
| Inducible HeLa LAP-DUSP7-56-181aa stable cell line | This paper | N/A |
| Inducible HeLa LAP-DUSP7-252-384aa stable cell line | This paper | N/A |
| Inducible HeLa LAP-DUSP7-252-419aa stable cell line | This paper | N/A |
| Inducible HeLa LAP-ERK2 stable cell line | This paper | N/A |
| Inducible HeLa LAP-ERK2-2A (T185A/Y187A) stable cell line | This paper | N/A |
| Inducible HeLa LAP-ERK2-2D (T185D/Y187D) stable cell line | This paper | N/A |
| Inducible HeLa LAP-ERK2-2E (T185E/Y187A) stable cell line | This paper | N/A |
| Oligonucleotides | | |
| siRNA targeting DUSP7 | Thermo Fisher Scientific | Cat# 4390824; siRNA ID: s4381 |
| Primer1 for DUSP7 qPCR: Fwd 5'-GACGTGCTCGGCAAGTATG-3' | Eurofins Genomics | N/A |
| Primer1 for DUSP7 qPCR: Rev 5'-GGATCTGCTTGTAGGTGAACTC-3' | Eurofins Genomics | N/A |
| Primer2 for qPCR: Fwd 5'-TGTGGCCTATCTGATGCAGAA-3' | Eurofins Genomics | N/A |
| Primer2 for qPCR: Rev 5'-GGGCGAGATGTTGGACTTTTTC-3' | Eurofins Genomics | N/A |
| Primer for cloning DUSP7-C331A: Fwd 5'-TGATGCCTGCCAGGGCGTGCACCAGGACAC-3' | Eurofins Genomics | N/A |

| | | |
|---|---|---|
| Primer for cloning DUSP7-C331A: Rev 5'-GTGTCCTGGTGCACGCCCTGGCAGGCATCA-3' | Eurofins Genomics | N/A |
| Primer for cloning DUSP7-R337A: Fwd 5'-GACCGTCACTGAGGCGCTGATGCCTGCC-3' | Eurofins Genomics | N/A |
| Primer for cloning DUSP7-R337A: Rev 5'-GGCAGGCATCAGCGCCTCAGTGACGGTC-3' | Eurofins Genomics | N/A |
| Primer for cloning DUSP7-1-181aa: Fwd 5'-GGGGACAAGTTTGTACAAAAAAGCAGGCTTCATGGGGAAAAACCAGCTCCGCGGCCCCCCAGCG-3' | Eurofins Genomics | N/A |
| Primer for cloning DUSP7-1-181aa: Rev 5'-GGGGACCACTTTGTACAAGAAAGCTGGGTCTCACTCTGTTTGAAACTTGTTG-3' | Eurofins Genomics | N/A |
| Primer for cloning DUSP7-56-181aa: Fwd 5'-GGGGACAAGTTTGTACAAAAAAGCAGGCTTCATGGGGAGCGCCGAGTGGCTGCAGG-3' | Eurofins Genomics | N/A |
| Primer for cloning DUSP7-56-181aa: Rev 5'-GGGGACCACTTTGTACAAGAAAGCTGGGTCTCACTCTGTTTGAAACTTGTTG-3' | Eurofins Genomics | N/A |
| Primer for cloning DUSP7-252-384aa: Fwd 5'-GGGGACAAGTTTGTACAAAAAAGCAGGCTTCATGGGGCTCTACCTCGGCTGCGCCAAGG-3' | Eurofins Genomics | N/A |
| Primer for cloning DUSP7-252-384aa: Rev 5'-GGGGACCACTTTGTACAAGAAAGCTGGGTCTCACGTCCGCTCAAAGTCCAGC-3' | Eurofins Genomics | N/A |
| Primer for cloning DUSP7-252-419aa: Fwd 5'-GGGGACAAGTTTGTACAAAAAAGCAGGCTTCATGGGGCTCTACCTCGGCTGCGCCAAGG-3' | Eurofins Genomics | N/A |
| Primer for cloning DUSP7-252-419aa: Rev 5'-GGGGACCACTTTGTACAAGAAAGCTGGGTCTCACGTGGACTCCAGCGTATTGAGTGG-3' | Eurofins Genomics | N/A |
| Primer for cloning ERK2-T185A/Y187A: Fwd 5'-TACCAACGTGTGGCCACAGCTTCTGCCAGGAACCCTGTGTGATC-3' | Eurofins Genomics | N/A |

| | | |
|---|---|---|
| Primer for cloning ERK2-T185A/Y187A: Rev 5'-GATCACACAGGGTTCCTGGCAGAAGCTGTGGCCACACGTTGGTA-3' | Eurofins Genomics | N/A |
| Primer for cloning ERK2-T185D/Y187D: Fwd 5'-TGTACCAACGTGTGGCCACATCTTCATCCAGGAACCCTGTGTGATCATG-3' | Eurofins Genomics | N/A |
| Primer for cloning ERK2-T185D/Y187D: Rev 5'-CATGATCACACAGGGTTCCTGGATGAAGATGTGGCCACACGTTGGTACA-3' | Eurofins Genomics | N/A |
| Primer for cloning ERK2-T185E/Y187E: Fwd 5'-CCCTGTACCAACGTGTGGCCACCTCTTCCTCCAGGAACCCTGTGTGATCATGG-3' | Eurofins Genomics | N/A |
| Primer for cloning ERK2-T185E/Y187E: Rev 5'-CCATGATCACACAGGGTTCCTGGAGGAAGAGGTGGCCACACGTTGGTACAGGG-3' | Eurofins Genomics | N/A |
| Recombinant DNA | | |
| DUSP7 cDNA | GenScript | Clone ID: OHu03759C; NM_001947 |
| pDONR221-MAPK1 (original clone FLH182003.01X) | DNASU Plasmid Repository | Clone ID: HsCD00076104 |
| pGLAP1-DUSP7 | This paper | N/A |
| pGLAP1-DUSP7-C331A | This paper | N/A |
| pGLAP1-DUSP7-R337A | This paper | N/A |
| pGLAP1-DUSP7-1-181aa | This paper | N/A |
| pGLAP1-DUSP7-56-181aa | This paper | N/A |
| pGLAP1-DUSP7-252-384aa | This paper | N/A |
| pGLAP1-DUSP7-252-419aa | This paper | N/A |
| pCS2-HA-DUSP7 | This paper | N/A |
| pCS2-Flag-DUSP7 | This paper | N/A |
| pCS2-Flag-DUSP7-C331A | This paper | N/A |
| pCS2-Flag-DUSP7-R337A | This paper | N/A |
| pGBioID2-DUSP7 | This paper | N/A |
| pGLAP1-ERK2 | This paper | N/A |
| pGLAP1-ERK2-2A(T185A/Y187A) | This paper | N/A |
| pGLAP1-ERK2-2D(T185D/Y187D) | This paper | N/A |
| pGLAP1-ERK2-2E(T185E/Y187E) | This paper | N/A |
| pCS2-HA-ERK2 | This paper | N/A |
| pCS2-Flag-ERK2 | This paper | N/A |
| pCS2-Flag-ERK2-2A(T185A/Y187A) | This paper | N/A |
| pCS2-Flag-ERK2-2D(T185D/Y187D) | This paper | N/A |
| pCS2-Flag-ERK2-2E(T185E/Y187E) | This paper | N/A |
| Software and Algorithms | | |
| GraphPad Prism 5 | GraphPad | RRID: SCR_002798 |

| | | |
|---|---|---|
| Adobe Photoshop | Adobe | http://shop.adobe.com/store/adbehap/DisplayHomePage |
| ImageJ | NIH ImageJ | https://imagej.nih.gov/ij/index.html |
| Biorender | Biorender | RRID:SCR_018361 |

#### LEAD CONTACT AND MATERIALS AVAILABILITY

#### METHODS DETAILS

##### Cell Culture

HeLa cells were grown in DMEM/Ham's F-12 with L-Glutamine (Genesee Scientific, El Cajon, CA) supplemented with 10% FBS in 5% $CO_2$ at 37 °C. HCT116 cells were grown in McCoy's 5A (Modified) Medium (Thermo Fisher Scientific, Canoga Park, CA) supplemented with 10% FBS in 5% $CO_2$ at 37 °C. For G1/S arrest and release, cells were arrested with 2mM thymidine (Sigma-Aldrich, St. Louis, MO) for 16-18 hours and washed three times with PBS, two times with complete medium before being released into fresh medium. For mitotic arrest, cells were arrested with 100nM Taxol (Sigma-Aldrich, St. Louis, MO) or 330nM nocodazole (Sigma-Aldrich, St. Louis, MO) for 18 hours. For metaphase arrest, cells were arrested with 10μM MG132 (Millipore Sigma, Burlington, MA) for one hour post thymidine release. For MEK inhibition, cells were treated with 50μM U0126 (Selleckchem, Houston, TX) for two hours or 18 hours in the case of immunoprecipitation. For ERK2 inhibition, cells were treated with 50μM FR 180204 (Selleckchem, Houston, TX) for two hours. For siRNA experiments, cells were mock transfected as negative control or transfected with Silencer Select Validated siRNA from Thermo Fisher Scientific (Canoga Park, CA) targeting DUSP7 (siRNA ID:s4381, cat. no. 4390824) at 20μM using Lipofectamine RNAiMAX (Thermo Fisher Scientific, Canoga Park, CA) for 72 hours before harvesting or fixation.

##### Plasmids, Mutagenesis, and Generation of Stable Cell Lines

DUSP7 and ERK2 sites mutants were generated by QuikChange Lightning Site-Directed Mutagenesis Kit (Agilent, Santa Clara, CA). cDNAs of full length GFP/DUSP7/DUSP7-C331A/DUSP7-R337A/ERK2/ERK2-2A/ERK2-2D/ERK2-2E and DUSP7 truncations were cloned into the C terminus of either pGLAP1, pGBioID2, pCS2-HA or pCS2-Flag *via* Gateway LR Clonase reaction (Torres et al., 2011). For transient transfection, HeLa cells were transfected with the plasmids generated above with FuGENE HD (Promega, Madison, WI) for 24-36 hours before harvesting or fixation. pGLAP1-only/DUSP7/DUSP7-C331A/DUSP7-R337A/ERK2/ERK2-2A/ERK2-2D/ERK2-2E/DUSP7-truncations and pGBioID2-only/DUSP7 were used to generate doxycycline inducible HeLa Flp-In T-REx LAP-GFP/DUSP7-C331A/DUSP7-R337A/ERK2/ERK2-2A/ERK2-2D/ERK2-2E/DUSP7-truncations and HeLa Flp-In T-REx BioID2-only/DUSP7 stable cell lines as described previously (Bradley et al., 2016; Torres et al., 2009). Briefly, pGLAP1- and pGBioID2-tagged genes of interest were co-transfected with pOG44 Flp-Recombinase Expression Vector (Thermo Fisher Scientific, Canoga Park, CA) into HeLa Flp-In T-REx cell lines by FuGENE 6 (Promega, Madison, WI). After selecting the integrated cells with 400μg/ml Hygromycin B (Thermo Fisher Scientific, Canoga Park, CA), the individual colonies were collected and continually grown for protein induction test before further protein purification experiments..

**LAP/BioID2 Purifications and LC-MS/MS Analyses**

For LAP purifications, GFP as the negative control and DUSP7 were purified from Taxol arrested LAP-tagged inducible stable cell lines as previously described (Torres et al., 2009). Briefly, LAP-only and LAP-DUSP7 stable cell lines were induced with 0.1μg/ml doxycycline (Sigma-Aldrich, St. Louis, MO) and treated with 100nM Taxol for 18 hours before being harvested and lysed. The cell lysates were subjected to tandem affinity purification by incubating with anti-GFP antibody beads; the bound eluates were incubated with S-protein Agarose (Millipore Sigma, Burlington, MA). The final eluates were resolved on a 4-20% gradient SDS PAGE gel (Bio-Rad, Hercules, CA); the gel was excised and prepared for LC-MS/MS analysis. For BioID2 purifications, biotinylated proteins were purified from Taxol arrested BioID2-tagged inducible stable cell lines using protocols described previously with modifications (Gupta et al., 2015; Kim et al., 2016). Briefly, BioID2-only and BioID2-DUSP7 stable cell lines were washed with PBS and DMEM/Ham's F-12 before being shifted into DMEM/Ham's F-12 supplemented with 10% Dynabeads (Thermo Fisher Scientific, Canoga Park, CA) treated FBS (FBS was incubated with Dynabeads at 4 °C overnight and the Dynabeads were removed with magnetic stand the following day). The cells were induced with 0.1μg/ml doxycycline and treated with 100nM Taxol and 50 μM Biotin (Sigma-Aldrich, St. Louis, MO) for 16 hours before being lysed in lysis buffer (50 mM Tris-HCl pH 7.5, 150 mM NaCl, 1 mM EDTA, 1 mM EGTA, 1% Triton-X-100, 0.1% SDS, protease inhibitor cocktail) for 1 hour at 4 °C with gentle rotation. The cell lysates were centrifuged at 15,000 rpm for 15 minutes and transferred to TLA-100.3 tubes (Beckman Coulter, Indianapolis, IN) for a second high speed centrifuge at 45,000 rpm for 1 hour at 4° C. The supernatants were incubated with Dynabeads at 4 °C overnight with gentle rotation. The beads were washed twice with 2% SDS, one time with WB1 (0.1% sodium deoxycholate, 1% Triton X-100, 500 mM NaCl, 1 mM EDTA, 50 mM HEPES), one time with WB2 (250 mM LiCl, 0.5% deoxycholate, 1 mM EDTA, 10 mM Tris-HCl pH 8.0), and a final wash with 50 mM Tris-HCl pH 7.5 before being resuspended in elution buffer (50 mM triethylammonium bicarbonate, 12 mM sodium lauroyl sarcosine, 0.5% sodium deoxycholate). The resuspended beads were proceeded to on-bead digestion and LC-MS/MS analysis. Mass spectrometry analysis was performed at the UCLA Pasarow Mass Spectrometry Laboratory on a Thermo LTQ-Orbitrap XL as described previously (Patananan et al., 2014). All raw mass spectrometry data were deposited at the UCSD Center for Computational Mass Spectrometry MassIVE datasets (TBD) and can be accessed using login: Torres, password: mitosis1. The LC-MS/MS results were analyzed and visualized with CANVS (Clean Analyze Network Visualization Software) (in preparation for publication). Protein-protein interaction information was derived and integrated from the Biological General Repository for Interaction Datasets (BioGRID v. 3.5) (Stark et al., 2006). Protein-complex information was derived from the Comprehensive Resource of Mammalian Protein Complexes (CORUM v. 3.0) (Giurgiu et al., 2019). Selected Gene Ontology (GO) terms (Gene Ontology release June 2019) (Ashburner et al., 2000) were used to analyze the protein-protein interactions based on cellular mechanisms. Final visualized affinity-based and proximity-based networks were generated with RCytoscapeJS (Franz et al., 2016; Shannon et al., 2003). All in-house R scripts incorporated in CANVS can be freely obtained at GitHub (TBD).

**Immunoprecipitations, *In Vitro* Binding Assays and Immunoblot Analyses**

For immunoprecipitations, LAP-tagged inducible stable cell lines were expressed with 0.1μg/ml doxycycline (Sigma-Aldrich, St. Louis, MO) for 18 hours and lysed in LAP200 lysis buffer (50mM Hepes pH 7.4, 200mM KCl, 1mM EGTA, 1mM MgCl2, 10% glycerol) plus 0.05% NP-40, 0.5mM DTT and protease inhibitor cocktail (Thermo Fisher Scientific, Canoga Park, CA). The cell lysates were incubated with S-protein Agarose (Millipore Sigma, Burlington, MA) at 4 °C for two hours. S-protein Agarose was then washed three times with LAP100 buffer (50mM Hepes pH 7.4, 100mM KCl, 1mM EGTA, 1mM MgCl2, 10% glycerol) plus 0.05% NP-40 and 0.5mM DTT. For *in vitro* binding assays, Flag-tagged GFP/DUSP7/DUSP7-C331A/DUSP7-R337A/ERK2/ERK2-2A/ERK2-2D/ERK2-2E and HA-tagged DUSP7/ERK2 were expressed in a Quick Coupled Transcription/Translation System (Promega, Madison, WI), incubated together with Anti-FLAG M2 magnetic beads (Sigma-Aldrich, St. Louis, MO) at 4 °C for 1.5 hours. The bound beads were washed three times with LAP200 buffer plus 0.05% NP-40 and 0.5mM DTT. The samples were resolved on a 4-20% gradient SDS PAGE gel (Bio-Rad, Hercules, CA) and transferred to a PVDF membrane (EMD Millipore, Burlington, MA). Membranes were incubated with primary antibodies in blocking buffer (PBS, 0.5% BSA, 0.05% Tween-20, 0.02% SDS, 0.05% Proclin) at 4 °C overnight, then washed three times (five-minutes each) with PBST (PBS, 0.1% Tween-20), and incubated with secondary antibodies conjugated to IRDye 680RD or IRDye 800CW at room temperature for 30 minutes. Western blots were scanned and analyzed on a LI-COR Odyssey Imager (LI-COR Biotechnology, Lincoln, NE). The relative intensity of western blot bands were analyzed with ImageJ (NIH, Bethesda, MD).

**Immunofluorescence and Live-cell Time-lapse Microscopy**

For immunofluorescence microscopy, HeLa cells were fixed with 4% paraformaldehyde, permeabilized with 0.2% Triton X-100/PBS, and blocked with IF buffer (PBS, 5% Fish Gelatin, 0.1% TritonX-100) before being incubated with 0.5 mg/ml Hoechst 33342 and the indicated primary antibodies in IF buffer at room temperature for one hour. Cells were then washed with PBS three times (five-minutes each) and incubated with secondary antibodies in IF buffer for 30 minutes. After a final wash, the coverslips were mounted with ProLong Gold Antifade mounting solution (Invitrogen, Carlsbad, CA) on glass slides. Images were captured with a Leica DMI6000 microscope (Leica DFC360 FX Camera, 63x/1.40-0.60 NA oil objective, Leica AF6000 software). Images were deconvolved with Leica Application Suite 3D Deconvolution software and exported as TIFF files. For quantification of defective cells, data represent the average ± SD of three independent experiments, with 100 cells counted for each. For live-cell time-lapse microscopy, HCT116 cells constitutively expressing GFP-H2B were imaged live six hours post thymidine release for 18 hours with an ImageXpress XL imaging system (Molecular Devices, San Jose, CA) at 37 °C in 5% $CO_2$ using a 20x air objective. Images were captured every five minutes with both bright field and FITC channels, and converted to AVI movies with ImageJ at one frame per second (NIH, Bethesda, MD).

**RT-qPCR**

Total RNA from mock transfected or DUSP7 siRNA transfected HeLa cells and DUSP7 cell lines were isolated with Direct-zol RNA Miniprep Kits (Zymo Research, Irvine, CA) and reverse transcribed with UltraScript 2.0 cDNA Synthesis Kit (Genesee Scientific, El Cajon, CA). qPCR was carried out with cDNA synthesized from above, Oligo(dT) Primers, and qPCRBIO SyGreen Blue Mix Lo-ROX (Genesee Scientific, El Cajon, CA) using a CFX Connect Real-Time PCR Detection System (Bio-Rad, Hercules, CA). qPCR data were analyzed with the Livak-Schmittgen method ($2^{-\Delta\Delta Cq}$) (Livak and Schmittgen, 2001).

**Antibodies**

Primary antibodies used for immunoprecipitations and immunofluorescence microscopy include: mouse anti-ERK2 (Santa Cruz Biotechnology, Santa Cruz, CA), chicken anti-GFP (Abcam, Cambridge, MA), rabbit anti-HA (Proteintech, Rosemont, IL), mouse anti-Flag (Sigma-Aldrich, St. Louis, MO), anti-FLAG Dylight 800 Conjugated (Rockland Immunochemicals, Limerick, PA), rabbit anti-pERK (CST, Denvers, MA), rabbit anti-pERK (R&D Systems, Inc., Minneapolis, MN), mouse anti-GAPDH (Proteintech, Rosemont, IL), rabbit anti-DUSP7 (Proteintech, Rosemont, IL), goat anti-Mad2 (Santa Cruz Biotechnology, Santa Cruz, CA), human anti-CREST (Antibodies Incorporated, Davis, CA), rat anti-α-tubulin (Bio-Rad, Hercules, CA), mouse anti-pERK (Abcam, Cambridge, MA), mouse anti-pERK (Santa Cruz Biotechnology, Santa Cruz, CA). Secondary antibodies conjugated to FITC, Cy3, and Cy5 were from Jackson ImmunoResearch Laboratories (West Grove, PA) and those conjugated to IRDye 680RD and IRDye 800CW were from LI-COR Biosciences (Lincoln, NE).

**QUANTIFICATION AND STATISTICAL ANALYSIS**

The replicate information and number of cells counted for each quantification were indicated in the figure legends. The data were analyzed using unpaired Student's t test in Figures 2D, 2G, 2K, 3C and 3F; paired Student's t test in Figures S2B and S2C. The Livak-Schmittgen method ($2^{-\Delta\Delta Cq}$) was used to analyze qPCR data. Data is judged to be statistically significant when $p< 0.05$. More specifically, asterisks indicate statistical significance as * $p< 0.05$, ** $p< 0.01$, *** $p< 0.001$. All statistical data were presented as the average±SD. All statistical figures were generated with GraphPad Prism 5.


Ashburner, M., Ball, C.A., Blake, J.A., Botstein, D., Butler, H., Cherry, J.M., Davis, A.P., Dolinski, K., Dwight, S.S., Eppig, J.T., *et al.* (2000). Gene ontology: tool for the unification of biology. The Gene Ontology Consortium. Nat Genet *25*, 25-29.

Bradley, M., Ramirez, I., Cheung, K., Gholkar, A.A., and Torres, J.Z. (2016). Inducible LAP-tagged Stable Cell Lines for Investigating Protein Function, Spatiotemporal Localization and Protein Interaction Networks. Journal of visualized experiments : JoVE.

Franz, M., Lopes, C.T., Huck, G., Dong, Y., Sumer, O., and Bader, G.D. (2016). Cytoscape.js: a graph theory library for visualisation and analysis. Bioinformatics *32*, 309-311.

Giurgiu, M., Reinhard, J., Brauner, B., Dunger-Kaltenbach, I., Fobo, G., Frishman, G., Montrone, C., and Ruepp, A. (2019). CORUM: the comprehensive resource of mammalian protein complexes-2019. Nucleic acids research *47*, D559-D563.

Gupta, G.D., Coyaud, E., Goncalves, J., Mojarad, B.A., Liu, Y., Wu, Q., Gheiratmand, L., Comartin, D., Tkach, J.M., Cheung, S.W., *et al.* (2015). A Dynamic Protein Interaction Landscape of the Human Centrosome-Cilium Interface. Cell *163*, 1484-1499.

Kim, D.I., Jensen, S.C., Noble, K.A., Kc, B., Roux, K.H., Motamedchaboki, K., and Roux, K.J. (2016). An improved smaller biotin ligase for BioID proximity labeling. Molecular biology of the cell *27*, 1188-1196.

Livak, K.J., and Schmittgen, T.D. (2001). Analysis of relative gene expression data using real-time quantitative PCR and the 2(-Delta Delta C(T)) Method. Methods *25*, 402-408.

Patananan, A.N., Capri, J., Whitelegge, J.P., and Clarke, S.G. (2014). Non-repair pathways for minimizing protein isoaspartyl damage in the yeast Saccharomyces cerevisiae. The Journal of biological chemistry *289*, 16936-16953.

Shannon, P., Markiel, A., Ozier, O., Baliga, N.S., Wang, J.T., Ramage, D., Amin, N., Schwikowski, B., and Ideker, T. (2003). Cytoscape: a software environment for integrated models of biomolecular interaction networks. Genome Res *13*, 2498-2504.

Stark, C., Breitkreutz, B.J., Reguly, T., Boucher, L., Breitkreutz, A., and Tyers, M. (2006). BioGRID: a general repository for interaction datasets. Nucleic Acids Res *34*, D535-539.

Tighe, A., Johnson, V.L., and Taylor, S.S. (2004). Truncating APC mutations have dominant effects on proliferation, spindle checkpoint control, survival and chromosome stability. Journal of cell science *117*, 6339-6353.

Torres, J.Z., Miller, J.J., and Jackson, P.K. (2009). High-throughput generation of tagged stable cell lines for proteomic analysis. Proteomics *9*, 2888-2891.

Torres, J.Z., Summers, M.K., Peterson, D., Brauer, M.J., Lee, J., Senese, S., Gholkar, A.A., Lo, Y.C., Lei, X., Jung, K., *et al.* (2011). The STARD9/Kif16a kinesin associates with mitotic microtubules and regulates spindle pole assembly. Cell *147*, 1309-1323.
